# High short tandem repeat mutation recurrence in heat-stressed *Arabidopsis thaliana* lines

**DOI:** 10.1101/2025.01.24.634728

**Authors:** William B. Reinar, Anne Greulich, Vilde O. Lalun, Melinka A. Butenko, Kjetill S. Jakobsen

**Author notes:** Contributed equally (William B. Reinar, Anne Greulich). Senior author (Melinka A. Butenko, Kjetill S. Jakobsen).

## Abstract

The rise of the annual average temperature presents a strong selection pressure for many species. Sessile organisms, such as the plant *Arabidopsis thaliana* (*A. thaliana*), need to adapt to elevated temperatures. Here, we conducted a mutation accumulation experiment in *A. thaliana* under high temperature to monitor genetic changes over nine generations in three parallel lines compared to three lines under normal growth conditions. Whole-genome sequencing of every sample allowed us to accurately capture mutations in troublesome regions by utilizing the genotypes of both parent and offspring. Our approach revealed that high temperature treatment disproportionally increased the rate of insertions and deletions in dimeric short tandem repeats (STRs) compared to the rate of single nucleotide variants. Recurrent mutations appeared more often than expected, indicating a mutation bias along the genome sequence. Our study sheds light on the mutational landscape of a plant genome when exposed to high temperature stress and points to STRs as key genomic elements that rapidly increase the standing genetic variation available for natural selection.

## Introduction

In addition to phenotypic plasticity, the standing genetic variation in *A. thaliana* populations and *de novo* mutations can drive adaptation in response to changing climates (Fournier-Level et al., 2011, Wilczek et al., 2014, Exposito-Alonso et al., 2019). A change in climate may produce stressful environments for plants such as *A. thaliana*, but its effect on genome stability and *de novo* mutation is poorly understood. The interaction between stress and mutation rates could be an important factor in genome evolution under stressful conditions but it can only be studied directly by performing mutation accumulation (MA) experiments.

Lu et al. 2021 performed MA experiments in *A. thaliana* under control and high temperature (CT and HT) growth conditions and revealed that plants propagated at HT displayed elevated mutation rates compared to plants grown at CT (Lu_HT/CT_: 32/23°C). Belfield et al. 2021 grew plants at HT (Belfield_HT_: 29°C) and compared the HT mutation rate with MA experiments under CT (23°C) by Jiang et al. 2014. Many of the mutations detected by Belfield and Lu were insertions and deletions (indels) within or near repetitive sequences, referred to in this study as short tandem repeat (STR) variants. However, the discrepancy of the reported fold increase (FI) estimates for STR variants under HT (3.6 vs. 19-23) renders the true effect of HT on STR mutations unclear.

It has long been suspected that STR sites have a role in eukaryotic gene regulation (Gemayel et al. 2010) and their spatial clustering in proximity to genes across the entire eukaryote Tree of Life leaves support for this hypothesis (Reinar et al. 2024). Certain coding STRs have been shown to impact protein activity in *A. thaliana* and rice (Reinar et al. 2021, Reinar et al. 2023, He et al. 2024). Since STR mutations correlates to expression levels of nearby genes in *A. thaliana* (Reinar et al. 2021) and in other systems (Gymrek et al. 2016, Fotsing et al. 2019) the reported elevation of STR mutation rates under heat-stress is intriguing: It entails that HT induces STR mutations with potentially important functional consequences that natural selection may act upon. Such heritable shifts in gene expression resembles phase variable expression of bacterial genes where length variation of STRs can serve as on/off gene-switches to facilitate rapid adaptation (Zhou et al. 2013, Wang et al. 2021).

Here, we conducted a new MA experiment in *A. thaliana:* We exposed six parallel *A. thaliana* lines to HT (30°C) and CT (22°C) for nine generations. To mitigate a potential high false-negative rate of indels within repetitive sequence, i.e., in STRs, we sequenced each individual plant (54 in total) by whole-genome sequencing (WGS) and used a specialized STR-variant caller (Willems et al. 2017) to map the occurrence and fate of *de novo* STR mutations. In summary, our study shows that the mutation rates of STRs are intensified under HT (FI = 2.6) and mainly occur within annotated transposable elements (TEs) and TE fragments. We observed a high number of independently recurring mutations (i.e., sites that mutated and accumulated a variant independently in parallel lines) beyond the expected, which indicates either an extreme HT-induced mutation bias along the genome sequence or recurrent, positive selection for the *de novo* variants. Either way, our study suggests that STR mutations make a disproportional but spatially restricted contribution to the standing genetic variation in a population under heat stress.

## Results

### High temperature treatment induced mutations

To investigate the effect at the whole genome level of HT on mutations in *A. thaliana,* we performed MA experiments under two temperature regimes: Three lines of *A. thaliana* Colombia-0 (Col-0) plants were grown at 22°C (CT; L1-L3) and three lines were grown at 30°C (HT; L4-L6) for nine generations (G0-G9) (**Fig. 1A)**. The 54 samples were subjected to WGS and variant calling (*Methods*). After adjusting for deeper sequencing of generation 8 and 9 (*Methods*) the mean sequencing depths were 21.2 reads per STR site per sample and 18.8 reads per SNV site per sample (**S. Fig. 1**). We leveraged the study design to assess the confidence in the *de novo* homozygous mutations (**Fig. 1B**).

**Fig. 1.**
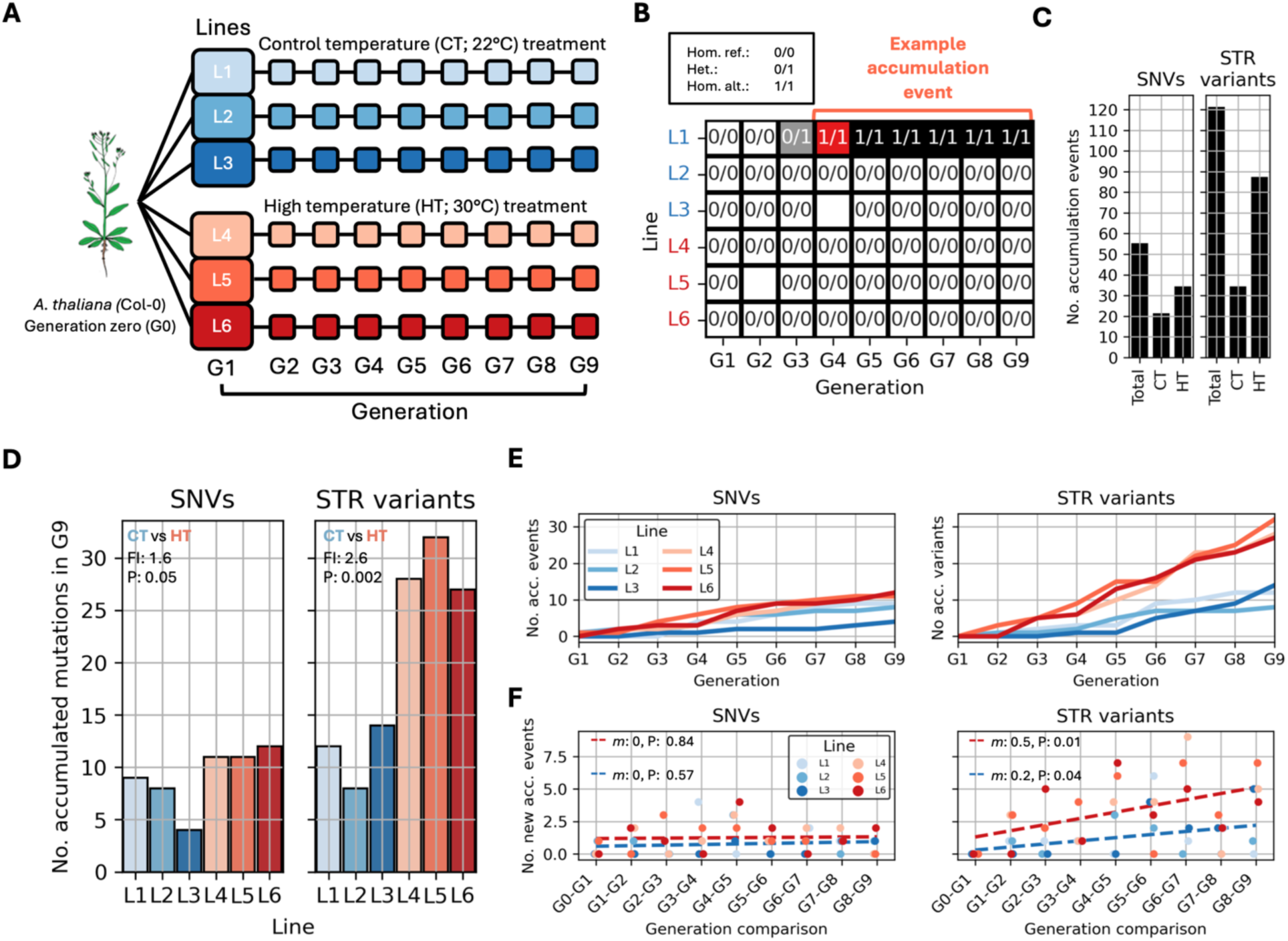
High temperature treatment induced mutations. **A.** The study design. Six lines (L1-L6) were grown over nine generations (ancestor G0 and G1-G9) at HT (30°C) and control temperature (CT; 22°C). Each sample was subjected to whole-genome sequencing. In all figures the CT lines L1-L3 are shaded in blue and the HT lines L4-L6 are shaded in red. **B.** An example accumulation event (at genomic position 3:13886242) where a homozygous reference genotype (Hom. ref.) mutated in G3 of L1 and appeared as a heterozygote (Het.), before segregating as a homozygote alternative genotype (Hom. alt.) in G4 and to the end of the experiment. White boxes without genotypes indicate missing genotypes. **C.** The number of accumulated STR variants and the number of accumulated single nucleotide variants (SNVs). **D.** The number of accumulated mutations in G9 per line (left: SNVs, right: STRs). The fold increase (FI) of the mean count of L4-L6 (HT) compared to the mean count of L1-L3 (CT) are indicated along with the P-value of an ANOVA test of equal means. **E.** The sum of accumulated SNV (left) and STR (right) variants per generation (x-axis; G1-G9) and per line. **F.** The number of new mutations (left: SNVs, right: STRs) per generation pair (G0-G1 – G8-G9). How well the data fits a linear function is denoted (*m* indicates the slope, and P indicates the P-value of the linear fit).

We counted 55 (HT: 34, CT: 21) SNV events and 121 (HT: 87, CT: 34) STR variant events yielding 0.5 SNVs per STR variant (**Fig. 1C**). As expected, HT treatment increased the occurrence of SNVs (SNV_CT_; L1-L3: 9, 8, 4, SNV_HT_; L4-L6: 11, 11, 12) and STR variants (STR_CT_; L1-L3: 12, 8, 14, STR_HT_; L4-L6: 28, 32, 27) yielding a mean HT-induced fold increase of 1.6 (SNVs) and 2.6 (STRs), but the fold change observed for SNVs due to HT treatment was not large enough to be statistically significant at the nominal P-value threshold (ANOVA_CT vs HT_ SNV_P-value_: 0.05, STR_P-value_: 0.002) (**Fig. 1D**).

Whereas SNVs accumulated at a steady rate (**Fig. 1E-F**), we observed that the number of new STR variants per generation increased over the course of the experiment **(Fig. 1E-F**): Given CT treatment, we observed an increase of 0.2 new STR variants per generation (linear fit_P-value_: 0.04) and an increase of 0.5 variants per generation under HT treatment (linear fit_P-value_: 0.01) (**Fig. 1F**). The SNV accumulation rate, however, did not vary significantly (linear fit: CT_P-value_; 0.57, HT_P-value_; 0.84) (**Fig. 1F**).

### High temperature treatment did not alter the mutation spectrum

Considering the SNV mutational spectrum of the 55 accumulated SNVs, the mean number of C|G (read C *or* G) → A|T mutations equaled 6.8 (± 2.2) per line, whereas the mean number of A|T → C|G equaled 2.0 (± 1.1) per line (ANOVA_P-value_ = 0.001), reflecting the C|G → A|T mutation bias known in many eukaryote and prokaryote systems (Lynch 2007, Hershberg and Petrov 2010). The pattern was consistent across the two treatments (CT_C|G→A|T L1-3_: 8, 6, 3 vs CT_A|T→C|G L1-L3_: 1, 2, 0, HT_C|G→A|T L4-6_: 10, 8, 6 vs HT_A|T→C|G L4-6_: 1, 2, 4) (CT_P-value_: 0.04, HT_P-value_: 0.02) **(Fig. 2A).** Considering STRs, we called 6,228 mononucleotide STR sites with motifs A|T of which only two sites accumulated a mutation (**Fig. 2B**). This number of mutated homopolymers was much lower than expected from literature, as Belfield reported almost 90 homopolymer variants during their experiment. Of the 3,408 AT|TA STR sites, 104 accumulated a mutation (**Fig. 2B**), which were higher compared to Belfield’s 24 dinucleotide repeat mutations (all but three with repeat motif AT|TA). The elevation of AT|TA mutation was present regardless of treatment (**Fig. 2C**). The number of insertions (62) almost equaled the number of deletions (59) (ratio: 1.05) and the ratios did not depend on treatment (CT_ratio_: 0.79, HT_ratio_: 1.18, Fisher’s exact test_P-value_: 0.41). The mean length of insertions (2.2 bp) was similar to the mean length of deletions (2.4 bp) (ANOVA_P-value: 0.12_) and similar in the CT and HT group (CT_mean length, insertions_:2.5 bp, CT_mean length, deletions_: 2.5 bp, ANOVA_P-value: 0.84_, HT_mean length, insertions_:2.1 bp, HT_mean length, deletions_: 2.4 bp, ANOVA_P-value: 0.06_) (**Fig. 2D**).

**Fig. 2.**
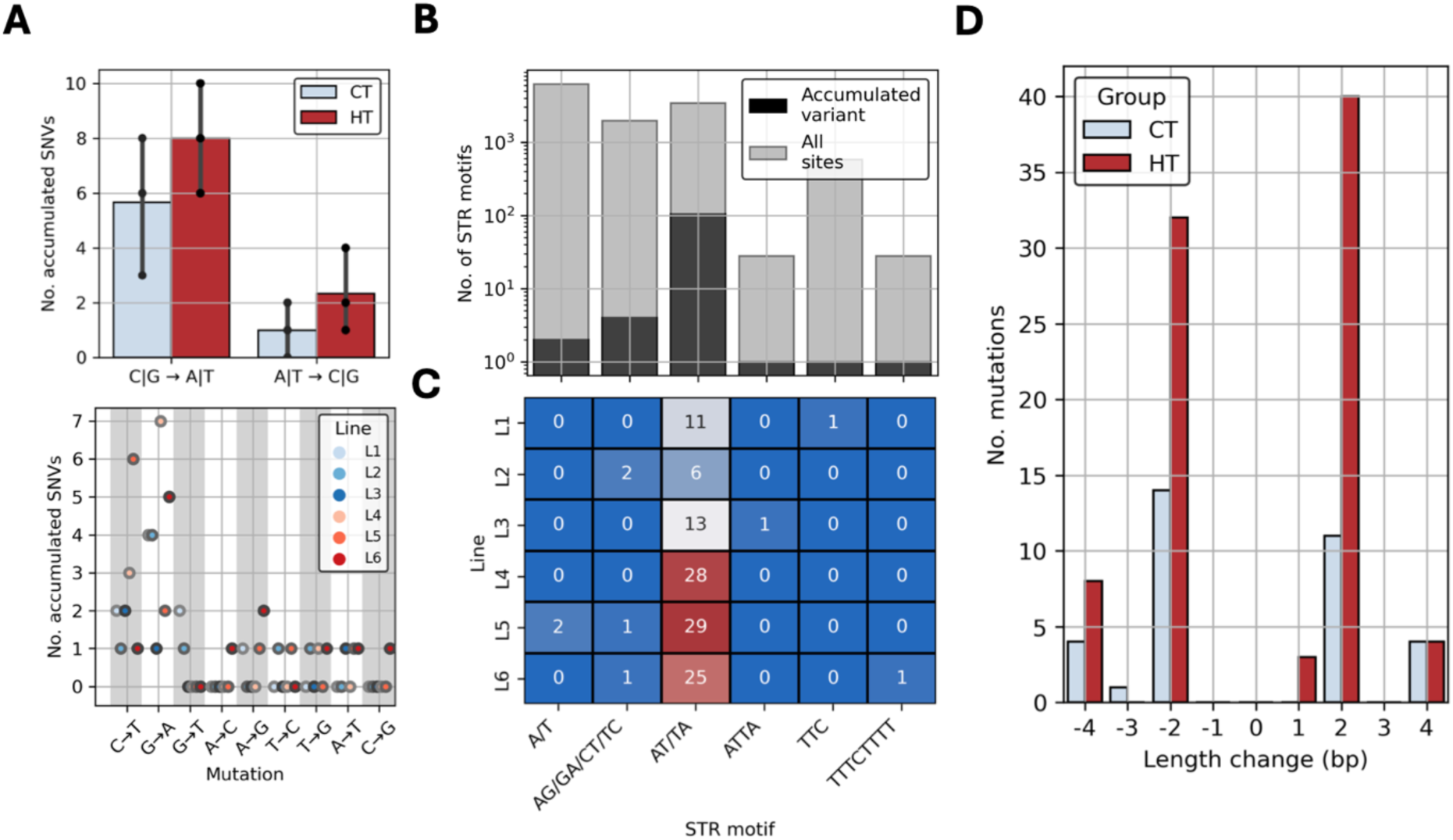
High temperature treatment did not alter the mutation spectrum. **A.** Top: The number of C|G → A|T (read C *or* G) and A|T → C|G single nucleotide variants (SNVs) conditioned on treatment group [HT; 30°C, control temperature (CT); 22°C]. Bottom: The number of SNVs in each line per specific transversion and transition. **B.** Gray bars show the number of STR motifs in the *Arabidopsis thaliana* Colombia-0 (The Arabidopsis Information Resource 10) reference genome. Black bars show the number of accumulated STR variants per motif. Note the log_10_-scaled y-axis. **C.** The heatmap indicates the number of STR sites with accumulated variants (cells shaded from blue to red according to the count) per motif (x-axis) and per line (y-axis). **D.** The histogram shows the length change (in bp, x-axis) of STR variants compared to the reference genotype, conditioned on HT and CT.

### GC-content correlates to short tandem repeat mutations across three studies

To investigate a relationship between mutation and GC-content we calculated the GC-content in 1 Kbp windows along the *A. thaliana* reference genome and correlated the window-specific GC-contents to the frequency of *de novo* mutations reported in Belfield et al. 2021, in Lu et al. 2021, and in this study. Across studies, STR variants were detected in windows with GC-contents exceeding 19% but below 45% (**Fig. 3A**). SNVs appeared in windows with GC-contents exceeding 21% but below 52% (**Fig. 3A)**. Within these ranges, the frequency of variants was partly explained by quadratic functions (STR_formula_: 2.5 x 10^-5^ x^2^ –0.002x + 0.05, R^2^: 0.84, P-value: < 0.001, SNV_formula_: 2.0 x 10^-5^ x^2^ – 0.001x + 0.03, R^2^: 0.21, P-value: 0.03) **(Fig. 3B-C**). Thus, local GC-content within these ranges is associated with the probability of *de novo* STR variants and SNVs (**Fig. 3B-C**). We investigated if the lack of SNVs in the tails of the genomic GC-distribution could be attributed to poor calling accuracy: We found that regions with GC-contents exceeding 52% and less than 21% could be partly attributed to lower calling accuracies in these regions as the proportion of variant calls passing hard-filtering with GATK was significantly lower (0.156) in regions exceeding 52% GC-content and lower (0.202) in regions with less than 21% GC-content, compared to the intermediate GC-range (0.268) (Fisher’s exact test P-values < 0.01) (**S. Fig. 2)**. However, the explanation may well be that the number of 1 Kbp windows at the tails of the genomic GC-distribution are diminishingly few (0.3% of the genome) and zero variants can thus be expected.

**Fig. 3.**
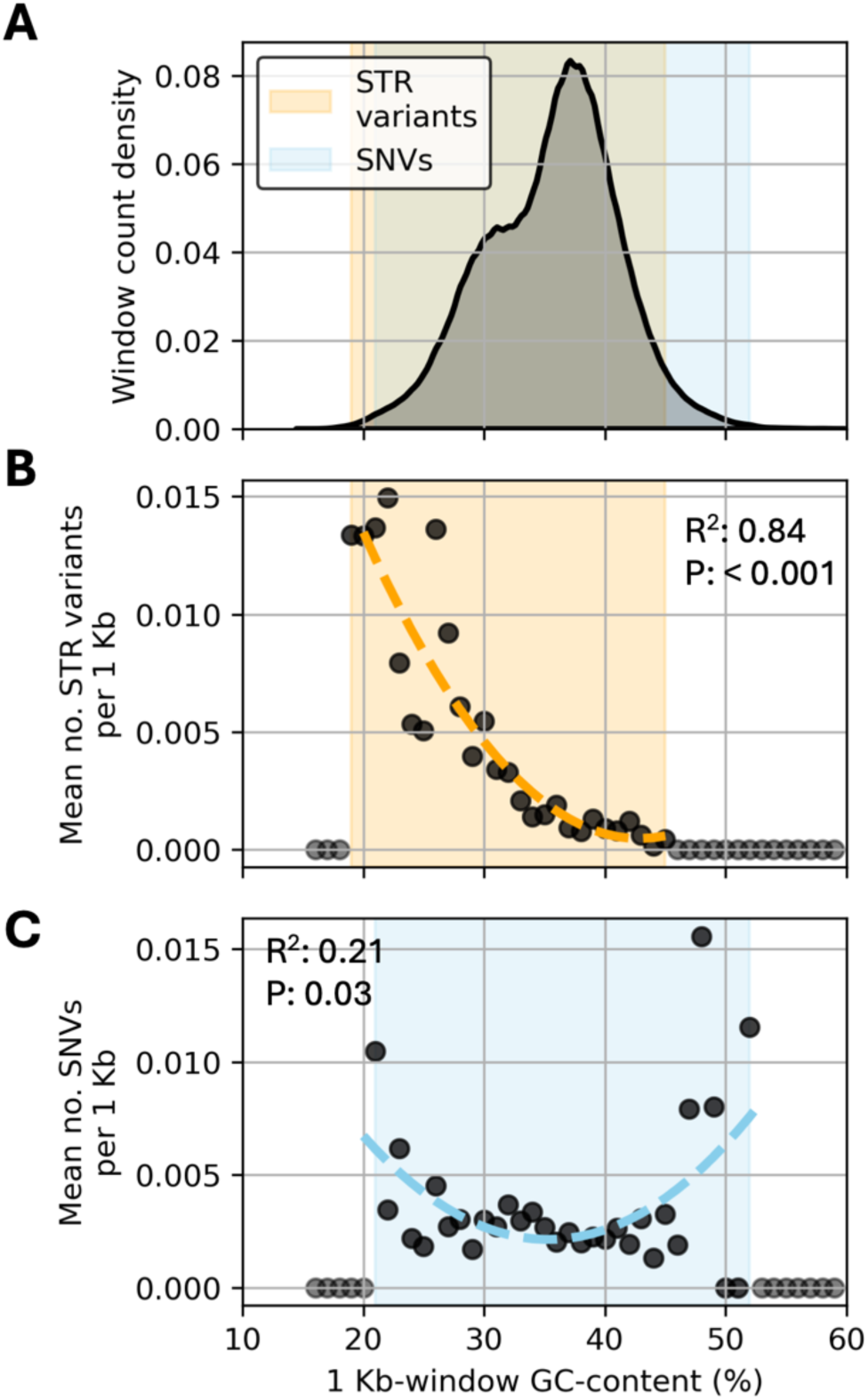
GC-content correlates to short tandem repeat mutations across three studies. Note the common x-axis for A, B, and C. **A.** The density plot shows the number of 1 kb windows (as density of counts, y-axis) with specific GC-contents (x-axis). The shaded light blue area shows the GC-content range where SNVs occurred across Belfield et al. 2021, Lu et al. 2021, and this study. The shaded light orange area shows where STR variants occurred across the three studies. The average number of STR variants (**B**) and single nucleotide variants (SNVs, **C**) from the three studies per 1 Kbp window as a function of the GC-content of the window. The lines show the fit of quadratic functions. The proportion of explained variance (R^2^) and the empirical P-value are denoted. The test was performed using data within the respective shaded areas (black, solid points).

### Mutation accumulation biased towards transposable elements across treatments

Of all called sites, 55,866 out of 145,682 sites were in annotated TEs (TE/non-TE ratio = 0.38), compared to 75 of the 168 sites that accumulated variants (TE/non-TE ratio = 0.81), which indicates a bias in mutation towards dispersed repeats versus unique sequences (Fisher’s exact P-value = 3.7 x 10^-6^) (**Fig. 4A**). TE-sequences accumulated STR variants beyond the expected (TE/non-TE ratio = 0.85, Fisher’s exact P-value = 9.5 x 10^-7^) and SNVs too displayed a higher, but non-significant TE/non-TE ratio (0.72, Fisher’s exact P-value = 0.053) (**Fig. 4A**). Accumulation of variants in TEs vs non-TEs did not depend on CT or HT treatment (SNV_CT vs HT Fisher’s exact P-value_ = 1.0, STR_Fisher’s exact P-value_ = 0.43 (**Fig. 4A**). We noted that some TE families accumulated more mutations compared to other TE families (**Fig. 4B**; left panel). Indeed, two TE families were statistically enriched for mutations, both under HT (BRODYAGA1A and HELITRONY3, Fisher’s Exact test P-values < 0.001) (**Fig. 4B**; right panel) despite that the number of accumulated variants in a TE family correlated to the number of elements of the TE family in the genome (linear fit R^2^ = 0.18, P-value = 0.002) (**Fig. 4B**; middle and right panel, **Fig. 4C**). The GC-content was slightly lower in genomic regions containing annotated TEs compared to the whole genome (GC_TEs_ = 34.5%, GC_whole-genome_ = 35.5%, T-test P-value < 0.001) and lower in regions containing mutated TEs compared to all TEs (GC_mut. TEs_ = 33.4%, P-value = 0.001) (**S. Fig. 3**). Together, these results show that the accumulated variants disproportionally occurred in TEs and that certain TE families accumulated more mutations than expected, as the number of accumulated mutations could not fully be explained by their overall counts in the genome nor by their, on average, lower GC-content, as the differences in GC-content was modest.

**Fig. 4.**
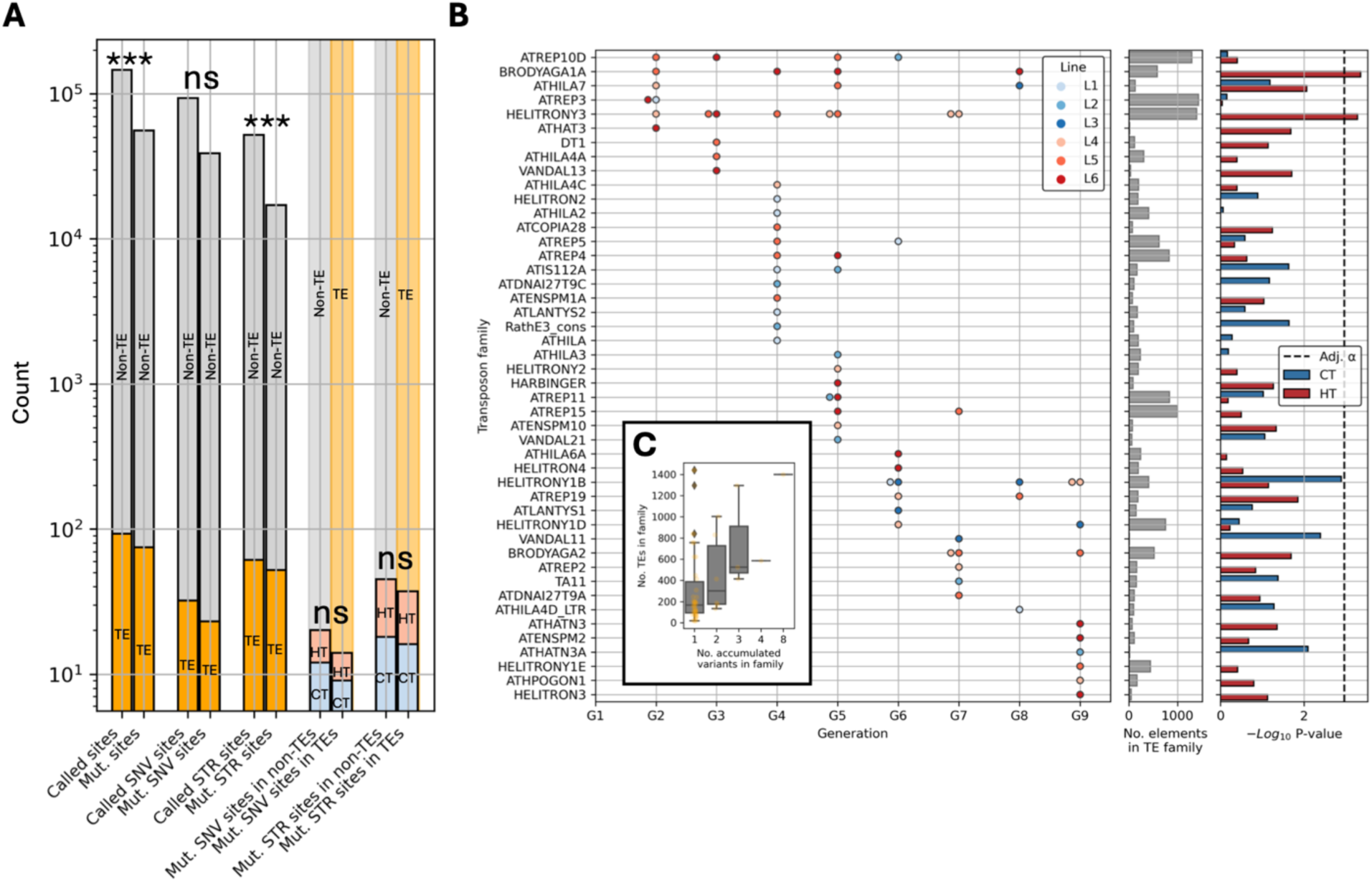
Mutation accumulation biased towards transposable elements across treatments. **A.** The bars show the number of sites overlapping transposable elements (orange) in the *Arabidopsis thaliana* The Arabidopsis Information Resource 10 (TAIR10) reference annotation. Gray bars indicate the number of non-overlapping sites (non-TE). The right-most bars indicate the number of sites with accumulated single nucleotide variant (SNV) or short tandem repeat (STR) variants in annotated transposable elements (orange background) and outside annotated transposable elements (gray background) conditioned on high temperature (HT; 30°C) and control temperature (CT; 22°C). ***, Fisher’s exact P < 0.001, ns, non-significant. **B.** Left: Accumulated SNVs and STR variants in annotated TE families (y-axis) per generation (G1-G9) and per line (CT lines L1-L3; shaded in blue, HT lines L4-L6; shaded in red). Middle: The bars show the number of TE elements in the TAIR10 reference genome of the given TE family (y-axis). Right: Enrichments of HT and CT variants in transposon families (one-sided Fisher’s exact test) **C.** The boxplots show the number of TEs in a family and are conditioned on the number of accumulated SNVs and STR variants in sequences deriving from the family (x-axis).

Although a disproportionate number of accumulated mutations occurred in TEs, 39 annotated genes accumulated STR or SNV mutations during the experiment. The accumulated genic mutations occurred in a subset of genes that did not deviate in function from the expected gene ontologies given the background set of gene ontologies (all False Discovery Rates > 0.05) **(S. Fig. 4**).

### More-than-expected recurrence of repeat mutations under high temperatures

No SNVs recurred among the lines. We did, however, observe seven sites with recurring STR mutations **(Fig. 5A**, **S. Fig. 5**). All the seven recurrent sites were in an annotated TE. None of the mutations recurred among CT lines (L1-3), whereas the HT lines (L4-L6) shared one or two mutations, and the CT and the HT lines shared three mutations (**Fig. 5B**). Even if the mutable scope was assumed limited to the sites where we observed a mutation (an extremely conservative assumption), observing zero overlap in the CT lines by chance still had a high probability (68.6%). Observing three shared mutations among CT and HT lines did not deviate strongly from the expected, as 10.0% of 1,000 simulations produced an overlap of at least three. However, observing five shared mutations within the HT lines had a probability of 2.3%, indicating that other processes than chance contributed to the recurrence of mutation under HT (**Fig. 5C**).

**Fig. 5.**
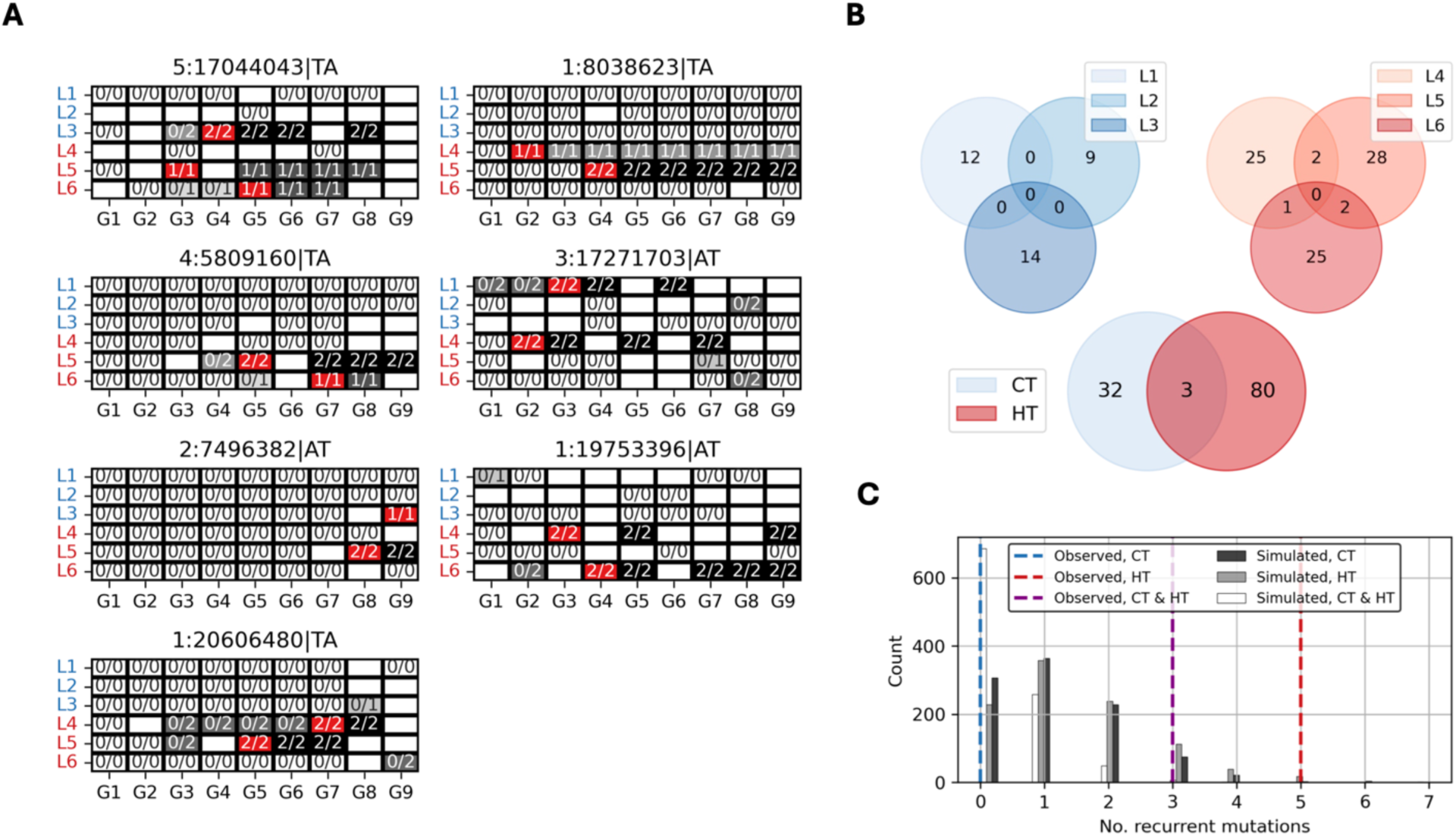
More-than-expected recurrence of repeat mutations under high temperatures. **A.** The eight sites with recurring mutation. The y-axes show the lines [control temperature (CT; 22°C) lines L1-L3 in blue, high temperature (HT; 30°C) lines L4-L6 in red] and the generations (G1-G9). Reference homozygous genotypes (0/0) are colored white, heterozygote genotypes (0/1) are colored light gray and homozygous alternative genotypes (1/1 or 2/2) are colored in black. White boxes without genotypes indicate missing genotypes. The generation in which the accumulated mutation first occurred is colored red. The genomic coordinate of the site in The Arabidopsis Information Resource 10 reference genome and the short tandem repeat (STR) motif are denoted in the title of each subplot. **B.** Venn diagrams showing the overlap of mutated STR sites within CT lines L1-L3 (top left), HT lines L4-L6 (top right), and CT lines and HT lines (bottom). **C.** The number of overlaps (x-axis) in 1,000 simulations given random placements of the observed accumulated STR mutation between CT lines (light blue), HT lines (red) and CT and HT lines (purple). The dashed vertical lines show the observed overlap within CT lines (blue), HT lines (red) and CT and HT (purple).

## Discussion

In this study we performed MA experiments in *A. thaliana* under CT and HT. The sequencing of all individual samples facilitated high-confidence inference of mutation in troublesome genomic regions (i.e., STR mutation within TEs that per definition are dispersed repeats). Our estimate of the HT-induced FI for SNVs (1.6) (**Fig. 1D**) was lower compared to reported estimates of 2-3 in the Belfield and Lu studies and our reported HT-induced FI of STR variants (2.6) (**Fig. 1D**) was lower than the FIs reported in Belfield and Lu (Lu_FI_ = 3.6, Belfield_FI_ = 19-23). A possible explanation for the discrepancies is that false-negative rates may be as high as 26% for STR variants (Belfield et al. 2021). Perhaps reflecting this high false-negative rate, the total number of STR variants differed dramatically between Lu and Belfield (57 vs 132), although the temperatures and the number of generations in the MA experiments were similar (10 generations at 32°C vs 11 generations at 29°C). The large discrepancy between Belfield’s reported HT-induced FI of 19-23 and this and Lu’s study likely stems from Belfield’s indirect comparison with CT-mutation estimates reported in earlier studies (Ossowski et al. 2010, Jiang et al. 2014) and does not conform to the HT-induced FIs revealed by direct comparisons, i.e., by identical sequencing and bioinformatics pipelines applied to control groups as in Lu et al. and in this study. Still, our data provides further evidence that HT treatment induces an increase in mutation rates (**Fig. 1D-E**).

The ratio of SNV/STR mutations (**Fig. 1C**) was similar in this study and in Belfield, whereas Lu found a higher SNV/STR ratio (0.5_This study_, 0.8_Belfield_, and 2.7_Lu_). Although it is not unconceivable that Col-0 founder plants maintained at different labs evolve different mutational spectrums, the discrepancy is maybe more likely be due to choices made in call filtering, which highlights the sensitivity to bioinformatics methodology in interpreting MA results. Our underestimation of homopolymer repeats compared to Belfield reflects a difference in methodology as well, since the specialized STR caller software (HipSTR) assigned all but a very few homopolymer variants as unreliable. Deeper sequencing and longer reads would probably alleviate the underestimation of homopolymer variants in our dataset.

The increase in the number of new STR mutations per generation, i.e., an increase of the STR mutation rate as the experiment progressed, was unexpected. The increase was almost twice as high under HT (*m*: 0.5) but non-negligible under CT *(m*: 0.2) (**Fig. 1F),** which may indicate that the CT-group was not completely stress-free, if stress was the root cause of the mutation rate increase. The increase could not be explained by differences in sequencing depth between lines and between generations as these were minimal (**S. Fig. 1**). A possible explanation for the successive increase in mutation rate under both treatments could be that the genetic bottleneck we introduced each generation led to cumulatively more genome instability leading to relaxed DNA-repair at mutation-prone STR sites. However, future replications of our MA study design (WGS of every generation or at regular intervals) in *A. thaliana* or in other species would be necessary to evaluate this hypothesis.

The SNV mutational spectrum was similar under CT and HT as transition and transversion ratios were similar across the treatments **(Fig. 2A**). AT/TA mutations dominated the STR mutational spectrum of both groups (**Fig. 2C**). As such, HT seemed to intensify but not substantially change the mutational spectrum. *A. thaliana* lines lacking DNA mismatch repair (MMR) displayed a ∼170 FI in SNVs and a ∼1000 FI in indel-mutations (Belfield et al. 2018). Mammalian studies have indicated that MMR is sensitive to stress (Chatterjee and Walker 2017), and if this is true also in *A. thaliana*, a stress-induced repression of MMR could explain the differences in the HT-induced FI we report for STR variants.

The high number of recurrently and independently mutated STR sites (seven) was not expected, although higher-than-expected parallel mutation has been reported in yeast (Kryazhimskiy et al. 2014). Independent, positive selection for the *de novo* variants cannot be ruled out as an explanation although it seems unlikely. The strong correlation between STR mutation and GC-content (R^2^ = 0.84, **Fig. 3B**) indicates that the sequence context is highly predictive of mutation in *A. thaliana*, consistent with empirical mutation models reported by Monroe et al. 2022. Considering this, the high mutation recurrence could perhaps be attributed to local mutation bias, which was intensified under HT treatment. Another possibility is that the recurrent mutations represent false positives but given the mutation trajectories where recurrent mutations occur as heterozygotes in an ancestor and are eventually fixed as homozygous in daughter and granddaughter plants, the recurrent genotypes seem very convincing (**Fig. 5A**). The additional independent lines with heterozygote variants at these mutation hotspots (**Fig. 5A**) would likely turn homozygous if the experiment had included more generations.

The U-shaped curve that could explain 21% of SNV occurrence across studies **(Fig. 3C)** indicates that the sites of point mutation are less predictable but still more likely to occur at the tails of the GC-content distribution. However, given the high number of possible point-mutable sites in the genome, a lack of recurring SNVs in 54 samples was expected.

The *A. thaliana* TE landscape mostly consists of old and thus degenerate elements not capable of transposition and make up approximately 20% of the *A. thaliana* genome sequence (Quesneville et al. 2020). Our results show that a disproportionate number of accumulated mutations, and all seven of the recurring mutations, occurred in these regions of the genome **(Fig. 4A**). This was true across treatments due to an overrepresentation of STR mutations, and not SNVs. Notably, specific TE families were enriched for mutation only in the HT group, suggesting that TE family-specific mutation rates may be influenced by HT (**Fig. 4B**).

In conclusion, our results show that HT treatment induced a substantial increase in STR mutation rates not observed for SNVs. By comparing STR mutations between treatment groups, we found that HT intensified the local genomic proneness towards STR mutation, which again depended on GC-content and whether the site was TE-derived. In total, 88 out of 121 variants we detected under HT were STR variants (**Fig. 1C**). If this pattern holds in natural populations exposed to stressful heat, it follows that 72% of the *de novo* mutations available for natural selection will be contributed by STRs, thus making up a substantial fraction of the new genetic variation that may be of importance under shifting environments.

## Resource availability

### Lead contact

Requests for further information and resources should be directed to and will be fulfilled by the lead contact, Kjetill S. Jakobsen unless otherwise noted.

### Materials availability

Seeds from *A. thaliana* lines grown in this study are available upon reasonable request through the senior author Melinka A. Butenko.

### Data and code availability

Raw reads from the samples sequenced in this study have been deposited to NCBI Short Read Archive at accession PRJEB70284. Code used for analysis, figures, and statistics are available at figshare (DOI: 10.6084/m9.figshare.28151285).

### Additional information

Any additional information required to reanalyze the data reported in this paper is available from the lead contact upon request.

## Acknowledgements

We thank Spyros Kollias, Morten Skage and A. Tooming-Klunderud from the Norwegian Sequencing Centre (NSC; https://www.sequencing.uio.no) for sequencing and processing the samples. Illumina library creation and high-throughput sequencing were carried out at the NSC, University of Oslo, Norway. We are grateful to Sergio Galino Trigo for helpful discussions. All computational analyses were performed on the Saga supercomputing cluster operated by UNINETT Sigma2, the National Infrastructure for High Performance Computing and Data Storage in Norway. The work was funded by Research Council of Norway (RCN) grant # 251076 (K.S.J and M.A.B) and by the Norwegian Ministry of Education and Research (fellowships to W.B.R, A.G, and V.O.L).

## Author contributions

K.S.J and M.A.B conceived the project. A.G performed experiments and bioinformatics. W.B.R performed bioinformatics and formal analyses. V.O.L provided expertise and feedback. W.B.R wrote the manuscript with input from A.G., V.O.L, M.A.B, and K.S.J.

## Methods

### Plant materials and growth conditions

Three *A. thaliana* lines grown at control temperature (CT, 22°C) and three *A. thaliana* lines grown at high temperature (HT, 30°C) by single seed descent for nine generations from the same wild-type *A. thaliana* Col-0 founder plant (G0) were established to investigate the effect of heat stress on STR length. For HT conditions a temperature of 30°C was chosen since plants grown at this temperature are known to exhibit temperature stress but still produce seeds (Whittle et al., 2009; Groot et al., 2017). Seeds were sown on MS-plates and stratified for 5 days at 4°C. Plates were transferred to Conviron growth chambers (16 h light/ 8 h dark) at constant 30°C or 22°C. Five days after germination plants were transferred to soil and grown to seed in the same growth chambers. Rosette leaf tissue was har­vested during or shortly after bolting.

### DNA extraction and whole-genome sequencing (WGS)

One *A. thaliana* plant of each generation and line (nine generations and three lines for each temperature, see **Fig. 1A**), as well as the founder plant of the experimental set-up, totaling 55 plants, were used for DNA extraction and WGS. Genomic DNA was extracted from rosette leaftissue using the ChargeSwitch™ gDNA Plant Kit (Invi­trogen). The DNA was sequenced using HiSeq4000 150-bp paired-end Illumina technology according to the manufacturer’s instructions at the Norwegian Sequencing Centre. Genera­tions 1 to 7 and generations 8 and 9 were sequenced in independent runs. This resulted in a difference in the average read depth (DP) for samples of the two runs. The average DP for generations 1 to 7 was 23.7 while for generation 8 and 9 aver­age DP was 45.3. To make samples more comparable, the binary alignment map (BAM) files of samples from generation 8 and 9 were subsampled to 23.7, the average DP of samples from generation 1 to 7. The subsampled data was used for all further analysis.

### Sequence alignment and variant calling

The reads of the sequencing data set were aligned to the *A. thaliana* TAIR10 reference genome (Lamesch et al. 2011; available from The Arabidopsis Information Resource at www.arabidopsis.org) using BWA (mem) (Li, 2013) and sorted using SAMtools (Li, 2009). The Spark implementation of Picard v.2.26.10 MarkDuplicates (https://github.com/broadinstitute/picard/) was used to mark duplicate reads in the BAM output files and remove them. Next, the TAIR10 reference genome was scanned for STRs using Tandem Repeats Finder (TRF) (Benson, 1999). The output of the TRF was used to generate an *A. thaliana* reference for the STR variant caller HipSTR described at https://github.com/HipSTR-Tool/HipSTR-references/. HipSTR v.0.6.2 (Willems et al., 2017) was used to call STR variants. HipSTR produced a Variant Call Format (VCF) output file used for further analysis. The BAM output files were used to call SNPs and indels using GATK v.4.2.0.0 HaplotypeCaller (Auwera and O’Connor, 2020). The gVCF output files were merged using CombineGVCFs and this merged gVCF file was genotyped using GenotypeGVCFs. The genotyped gVCF file was split into a SNP and indel file using GATK’s SelectVariants. Conforming to best practices the GATK SNV variants were initially filtered with the parameters QD < 2.0, QUAL < 30.0, SOR > 3.0, FS > 60.0, MQ < 59.5, MQRankSum < −12.5, and ReadPosRankSum < −8.0. Since our downstream analyses relied on a high confidence in G0 calls, G0 calls were required to be homozygous as we did not have sufficient read depth to accurately call heterozygotes. In addition, all SNV calls were required to have a GQ > 30. HipSTR variant calls were filtered using the recommended parameters (--min_call_qual 0.9, --max-call-flank-indel 0.15, --max-call-stutter 0.15, --min-call-allele-bias –2, and –min-call-strand-bias –2) as described at https://hipstr-tool.github.io/HipSTR/#call-filtering. Scikit-allel v.1.3.5 (Miles et al. 2023) and Pandas v.1.5.2 (McKinney, 2010) were used to load and parse the VCF files.

### Defining accumulated variants

The definition of an accumulated variant was based on information on the genotype in the parent and the offspring plant using a custom Python script (the function “getCalls” in the “Gen_Pipeline_and_Figures” Jupyter Notebook). A genotype was designated to be an accumulated homozygous variant if it met the following requirements: a homozygous genotype was 1) non-missing in G0, 2) not missing in the current generation, 3) not equal to the previous generation, 4) equal to all succeeding non-missing genotypes, and 5) not equal to G0. For the homozygous STR variants that first appeared in G9 (i.e., no data on succeeding genotypes) the genotypes from the raw HipSTR variant calling output was examined together with the read alignments of each generation in the Integrative Genome Browser web application (Robinson et al. 2011). Likewise, variants that appeared in a single sample but with missing high-quality calls in preceding and succeeding generations was assessed by the raw genotypes and read alignments of the preceding and succeeding generations. Further, read alignments were checked to identify duplicate calls of identical STR variants and to identify SNVs in STR variants.

### Meta-analysis of *de novo* mutations and GC-content

We parsed the coordinates of de novo variants and mutation types (indels and SNVs) from the supplementary data released with the Belfield study and the Lu study. These sites, and sites originating with this study, were used to build a BED file containing the genomic coordinates and the mutation type. To separate indels occurring in STR sites from indels in unique sequence we extracted the mutation site coordinate ± 10 bp from the *A. thaliana* TAIR10 reference genome and scanned the 10-bp sites for repetitiveness as in Reinar et al. 2024. Sequences with repetitiveness scores ≥ 0.5 (meaning that 50% of the 10 bases were repeated in tandem) were defined as STR sites if the mutation was designated as in indel in the original study. Next, we calculated the GC-content in 1 Kbp windows along the *A. thaliana* TAIR10 reference genome and formatted the results as a second BED file. These BED files were intersected using pybedtools v0.9.0 (Dale et al. 2011) to retrieve the genomic window and its associated GC-content and the number of STR variants and SNVs occurring in each window. In 1 Kbp windows with GC-contents less than 21%, within 21-52% and over 52% we counted the proportion of variants passing filtering based on the “FILTER_PASS” column in the hard-filtered GATK4.0 VCF. Two-sided Fisher’s exact tests as implemented in SciPy v.1.12.0 (Virtanen et al. 2020) were used to test for dependence between GC-content and the proportion of filtered variants.

### Intersections with TEs

We parsed the “TAIR10_Transposable_Elements.txt” file retrieved from www.arabidopsis.org to construct a BED file with TE genomic coordinates and TE family names. By using pybedtools v0.9.0, the TE coordinates were intersected with the coordinates containing the mutated sites reported in this study.

### Statistical analysis

#### Effect of treatment on the mean number of accumulated variants

A one-way ANOVA was used to test the hypothesis that the CT group (N = 3) had an equal number of accumulated homozygous mutations compared to the HT group (N = 3). As such, the mean of the three counts in the CT-group was compared with the mean of the three counts in the HT-group. The ANOVA-test was performed using the “f_oneway” function implemented in SciPy v.1.12.0.

#### Effect of generation on the number of new accumulated variants

To see if the number of new accumulated variants increased per generation as the experiment progressed, we tested if the linear fit of new accumulated variants as a function of the generation (G_1_ – G_2_, …, G_8_ – G_9_) had a slope deviating significantly from zero. The linear regression was performed the “linregress” function implemented in SciPy v.1.12.0.

#### Effect of treatment on mutational spectrum

A one-way ANOVA was used to test if the mean number of A|T → C|G mutations equaled the mean number of C|G → A|T mutations. The scope of the test was 1) all mutations (CT-group and HT-group), 2) within the CT-group, and 3) within the HT-group. The ANOVA-test was performed using the “f_oneway” function implemented in SciPy v.1.12.0. Equal insertion/deletion ratios were assessed using two-sided Fisher’s exact tests (“fisher_exact”) as implemented in SciPy v.1.12.0 specifically testing if the ratio of insertions vs deletions was equal among the CT and HT group. Equal insertion and deletion lengths were assessed by testing if the mean insertion length equaled the mean (absolute) deletion length using ANOVA as implemented in SciPy v.1.12.0. The scope of the tests was 1) all insertions and deletions, 2) insertions and deletions specific to the CT group, and 3) insertions and deletions specific to the HT group.

#### Quadratic fit of variant frequencies related to GC-content

The quadratic functions were estimated by the NumPy v.1.23.5 (Harris et al. 2020) “polyfit”-function inputting the GC-contents, the mean variant frequencies, and the degree of the fitting polynomial (second degree). After calculating the proportion of variance in variant frequencies that was explained by the quadratic functions (R^2^), we shuffled both GC-contents and variant frequencies 1,000 times and generated 1,000 new quadratic functions to produce a distribution of R^2^s. The number of R^2^s exceeding the observed R^2^ on the real data divided by the number of iterations (n = 1,000) produced an empirical P-value to indicate the statistical significance of the quadratic fit.

#### Effect of TEs on mutation

To assess if there was a dependence between sequence context (TE or non-TE sequence) and the number of accumulated variants we used two-sided Fisher’s exact tests as implemented in in SciPy v.1.12.0. The ratio of all called sites in TE-derived sequences and all called sites outside TE-derived sequences was compared to the ratio of TE-derived sites accumulating a mutation and non-TE sites accumulating a mutation. We performed the test for 1) all sites, 2) all SNV sites, and 3) all STR sites. We further tested, for STR variants and SNVs separately, if the TE/non-TE ratio depended on treatment (HT and CT) using two-sided Fisher’s exact tests. Using linear regression as implemented in the “linregress” function implemented in SciPy v.1.12.0 we regressed the number of TEs of a given TE family (based on information in the “TAIR10_Transposable_Elements.txt” file retrieved from www.arabidopsis.org) on the number of accumulated variants in the family (ranging from one to eight). Last, one-sided Fisher’s exact tests were used to test, separately for treatment groups, for dependence between TE family and a higher number of accumulated mutations. To correct for the number of tests we divided the nominal P-value threshold (0.05) by the number of elements with at least one fixed mutation.

#### Calculations of the number of expected recurrent mutations

After designating accumulated variants according to the procedure described in the Methods section “*Defining accumulated variants*” we generated a null expectation by defining an equal, but random, number of genotypes to be accumulated 1,000 times. We performed the procedure separately for the CT group and the HT group and recorded the overlap between lines within the CT group, within the HT group, and the overlap between accumulated (mock) genotypes in the CT group and the HT group. The probability of obtaining real counts given chance alone was estimated by counting the number of mock observations exceeding the real observation divided by the number of iterations (1,000).

#### Gene Ontology enrichment

Annotated TAIR10 gene IDs associated with genes overlapping mutation coordinates were queried in the PANTHER database *via* www.arabidopsis.org choosing Fisher’s exact test as the method for statistical analysis. P-values were adjusted using the Bonferroni method.

## Supplementary Figures

**S. Fig. 1.**
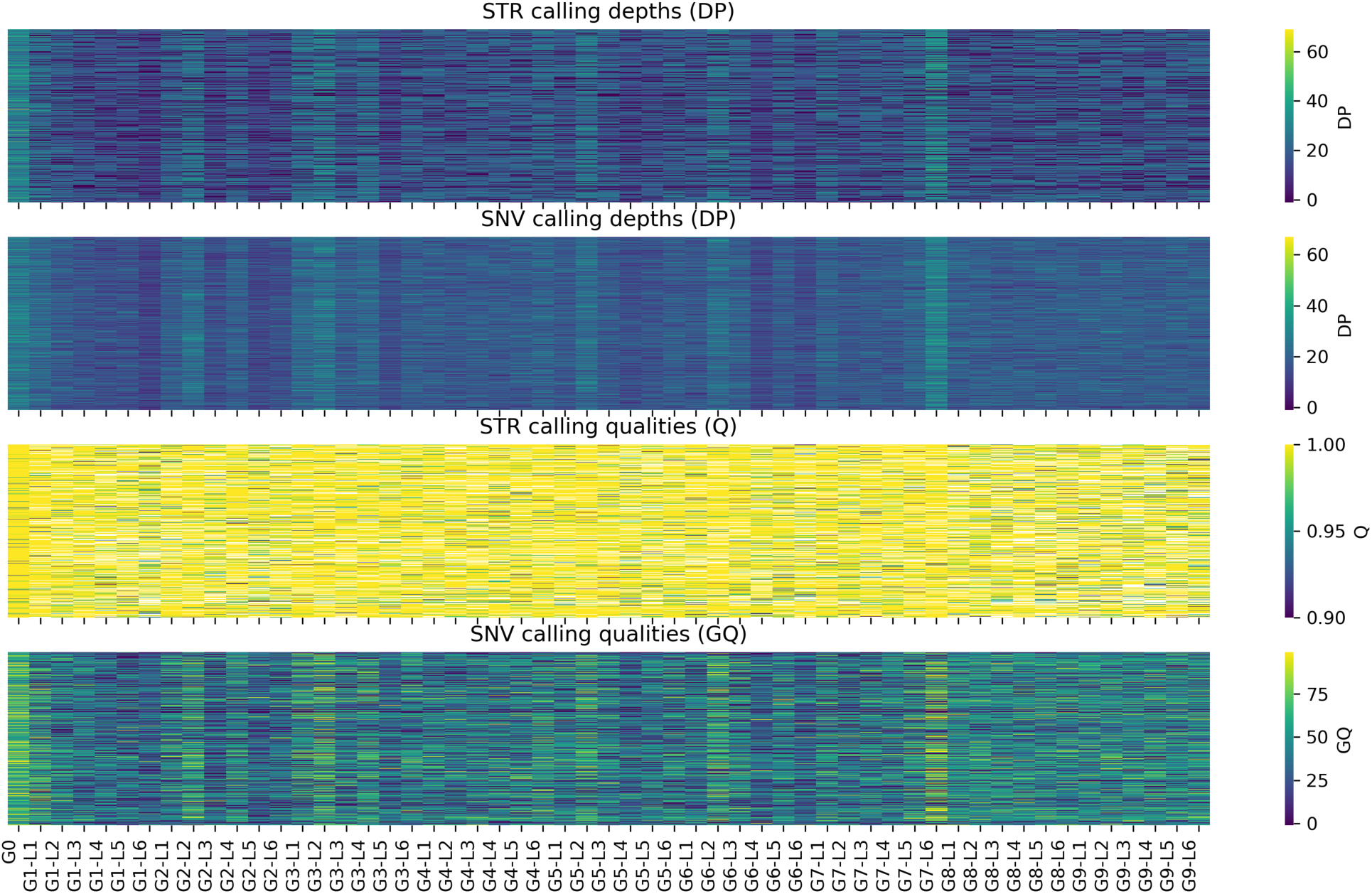
Sample depths and qualities after quality control. The heatmaps indicate the depths and qualities of short tandem repeat (STR) variant calls and single nucleotide variant (SNV) sites per sample. Note that G8-G9 appear to have very even sequencing depths and qualities since the reads of these generations were thinned after an initial deeper sequencing.

**S. Fig. 2.**
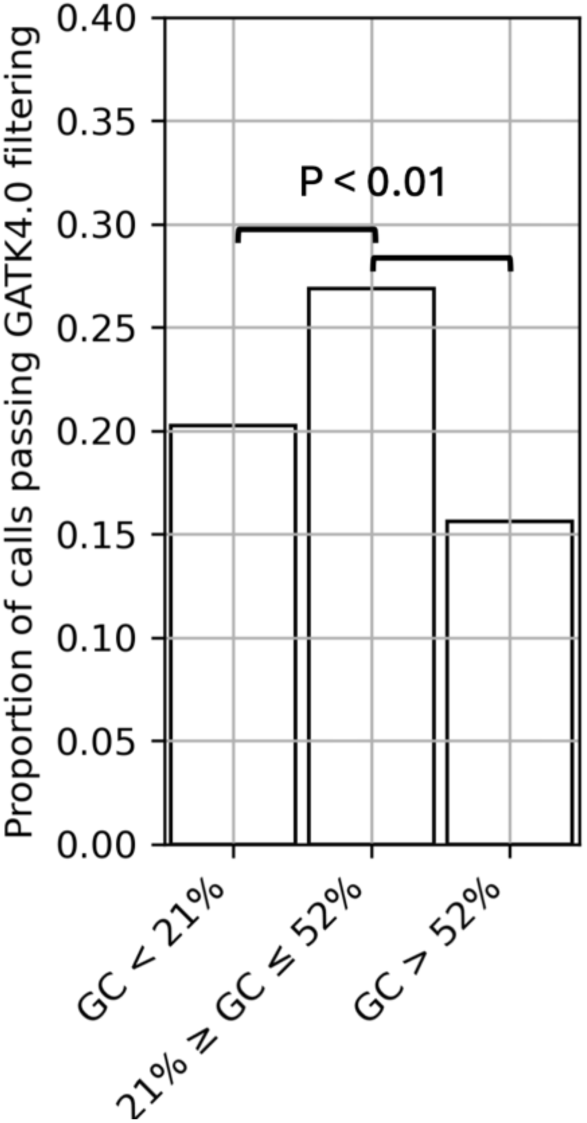
The proportion of calls passing filtering conditioned on GC-content. The y-axis indicates the proportion of variant calls that passed the GATK4.0 filtering (*Methods*) as a function of the GC-content (in 1 Kbp windows) containing the mutated site. The P-value indicates the result from two-sided Fisher’s exact tests.

**S. Fig. 3.**
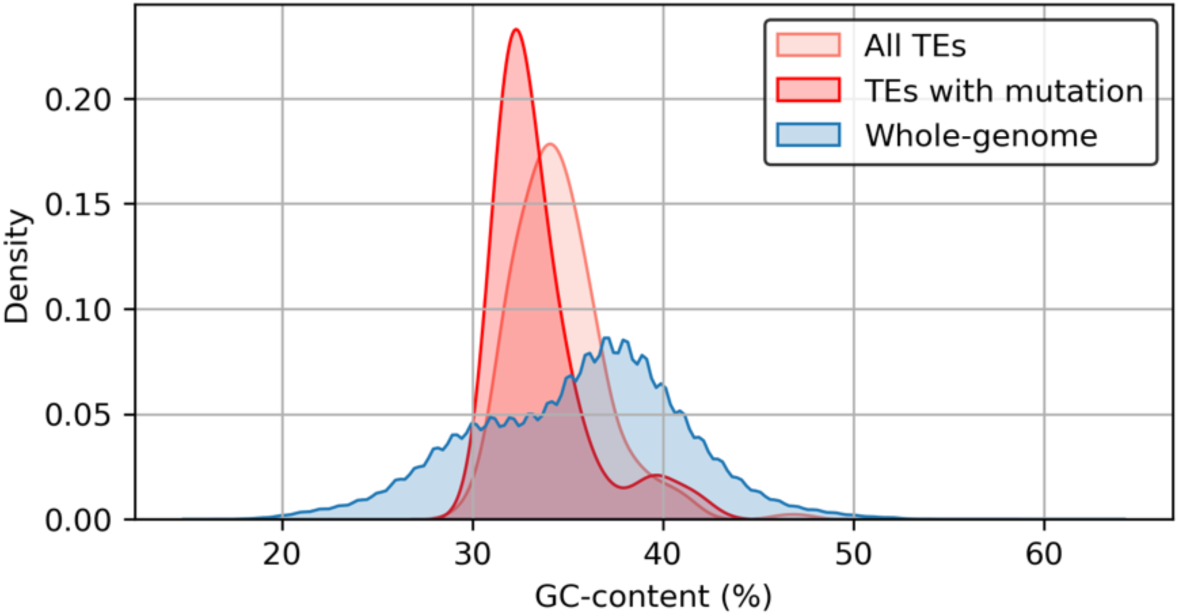
Distribution of GC-content along the genome and in transposable elements (TEs). The density curves indicate distributions of GC-content in 1 Kbp windows containing TEs (light red), TEs with mutation(s) (red), and 1 Kbp windows across the whole-genome (blue).

**S. Fig. 4.**
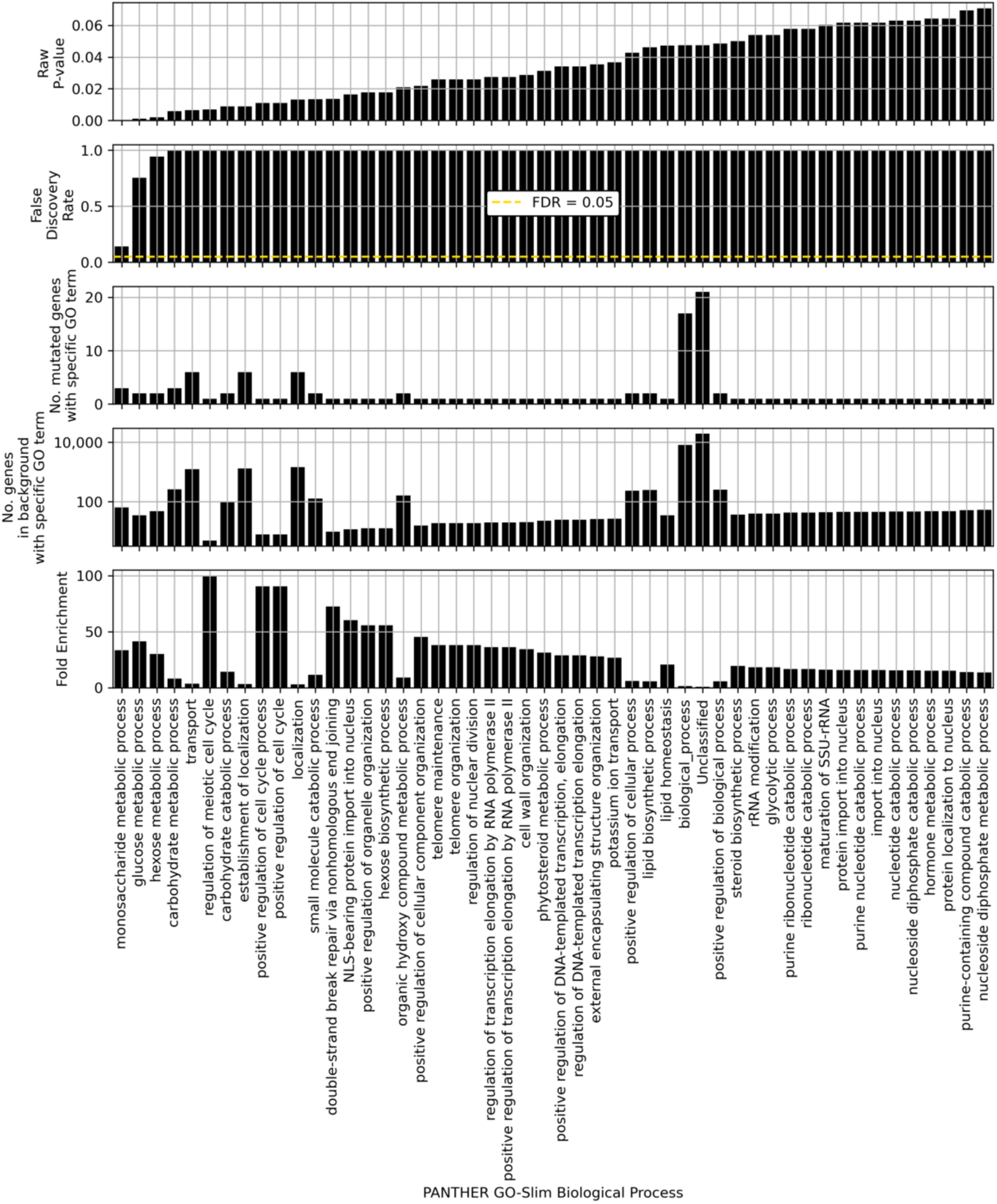
Gene Ontology (GO) biological processes of mutated genes compared to the *Arabidopsis thaliana* background set of GO biological process functions. All subpanels share the x-axis, which displays the top 50 PANTHER GO-Slim biological processes when sorted by raw P-values (top). The second panel indicates the False Discovery Rate (FDR) with the FDR = 0.05 denoted as a yellow, dashed line. The third panel shows the number of mutated genes per category (categories may overlap) and the fourth panel shows the number (log-scaled) of genes per category in the *Arabidopsis thaliana* TAIR10 background genes. The last panel shows the fold enrichment values when GO categories in the mutated gene list were compared to the background set of GO categories with one-sided Fisher’s exact tests.

**S. Fig. 5.**
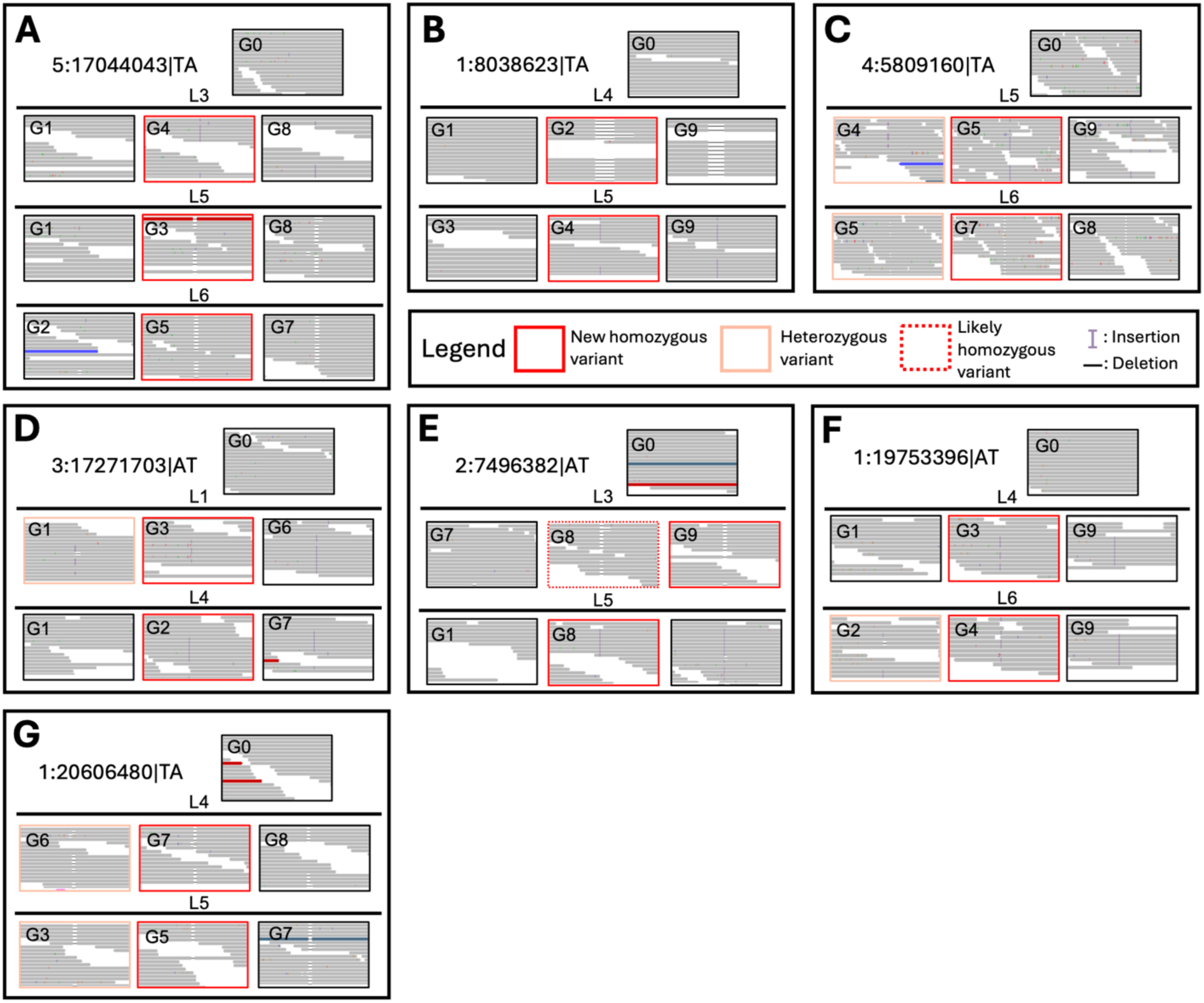
Read alignments of recurrent variants in short tandem repeats. **A-G** shows excerpts of the alignments of reads containing new homozygous insertions (red boxes), heterozygous variants (orange boxes), and likely false negative new homozygous insertions (dashed, red boxes). The alignments are shown for a selection of generations (G0-G9) and lines (L1-L6). Insertions are marked with a purple capital “I” in the alignment and deletions as a dash in the alignment. Reads colored blue in A and C indicates a lower-than-expected insert size and reads colored red in A, D, E, and G indicates higher-than-expected insert sizes. The dark blue colored reads in E and G indicates mapping of the reads’ mate pair to chromosome 5. Colored nucleotides indicate mismatches. The read alignments were visualized with the Integrative Genome Browser web application.

